# Is cold-induced nuclear import of CBF4 regulating freezing tolerance?

**DOI:** 10.1101/2022.08.11.503593

**Authors:** Wenjing Shi, Michael Riemann, Sophie-Marie Rieger, Peter Nick

**Author notes:** Author to whom correspondence should be addressed: Wenjing Shi, Molecular Cell Biology, Botanical Institute, Fritz-Haber-Weg 4, 76131 Karlsruhe, Germany.

## Abstract

C-repeat binding factors (CBFs) are crucial transcriptional activators in plant responses to low temperature. CBF4 differs by a slower, but more persistent regulation and its role in cold acclimation. Cold acclimation is of accentuated relevance for the tolerance to late spring frosts as they become progressively common as consequence of blurred seasonality in the context of global climate change. In the current study, we explore the functions of CBF4 from grapevine, VvCBF4. Overexpression of VvCBF4 fused to GFP in tobacco BY-2 cells confers cold tolerance. Furthermore, this protein shuttles from the cytoplasm to the nucleus in response to cold stress, associated with accumulation of transcripts for other CBFs and the cold responsive gene ERD10d. This response differs for chilling as compared to freezing and is regulated differently by upstream signalling involving oxidative burst, proteasome activity and jasmonate synthesis. This difference between chilling and freezing is also seen in the regulation of CBF4 transcripts in leaves from different grapevines differing in their cold tolerance. We propose the quality of cold stress is transduced by different upstream signals inducing nuclear import to regulate other CBF factor and activate COR genes.

## Introduction

Global warming, as one of the most fatal aspects of climate change, is influencing all the life stages of land plants. As revealed by a study on 500 plant taxa in Massachusetts (USA) and 400 British plant species, flowering has advanced since 1736 in average 4-7 days with each °C of increase, leading to precocious bud break driven by elevated temperature (Lippmann *et al*., 2019). Additionally, extreme weather phenomena like early autumn frost, late frost in spring, as well as cold and hot waves during the early summer occur more often, which leads to an aggravating negative impact on plants. For viticulture, a correlative analysis of harvest times and climatic conditions in France and Switzerland over the past four centuries showed that harvests have shifted to earlier time points, such that maturation is progressively uncoupled from the dry autumn weather required for high-quality wine (Cook and Wolkovich, 2016). A second problem is that the freezing tolerance of grapevine drops rapidly after bud break (Fuller and Telli, 1999). The progressively blurred onset of spring with unusually warm weather in March and late freeze episodes in April has, therefore, especially devastating consequences for viticulture. Thus, understanding the molecular mechanisms underlying cold stress and cold adaptation has turned into a very relevant research topic.

The quality of cold stress is strongly depending on the temperature and, traditionally, two versions of cold stress are discriminated (for classical reviews see (Lyons, 1973; Guy, 1990)): Chilling stress is defined as chilling temperatures above the freezing point (0-20°C), and affects productivity and quality of plants inhabiting tropical or subtropical climes (Zhang *et al*., 2020). Unlike chilling, the injury caused by freezing stress (< 0°C) is irreversible and can be lethal due to formation of intracellular ice crystals. Whether a given plant is tolerant to chilling or freezing, respectively, is strongly dependent on its genotype. In case of grapevine, *V. amurensis* can survive at −40°C, while severe cane damage of *Vitis vinifera* starts already from +2.5°C for Riesling and from −5°C for Pinot Noir, according to the USDA extension service (2021). The ability to respond adequately to different temperatures, obviously requires efficient and specific signalling.

In *Arabidopsis,* approximately 170 COR genes are CBF-dependent, involved in different adaptive responses, such as re-fluidisation of membranes, osmotic adjustment by compatible solutes, or protection of membranes and proteins by cryoprotective proteins and soluble sugars (Shi *et al*., 2018). The Arabidopsis homologues CBF1, CBF2, and CBF3, also known as DREB1b, DREB1c, and DREB1a–are rapidly induced in response to cold stress (Medina *et al*., 2011) and can, upon overexpression effectively activate downstream genes, thus enhancing their cold response (Zhou *et al*., 2017). Conversely, the *A. thaliana* null *cbf1 cbf2 cbf3 (cbfs) triple mutant* showed impaired chilling response and is extremely sensitive to freezing, even after a cold-acclimation treatment (Jia *et al*., 2016). A second tier of signalling allows not only for signal amplification, but also for signal diversification. In fact, loss-of-function mutants showed that *CBF1 and CBF3* are required for freezing tolerance as result of cold acclimation, while *CBF2* was dispensable in this respect and, thus, might convey alternative functions (Novillo *et al*., 2007). Furthermore, *AtCBF2* is expressed in a pattern that is different from *AtCBF1* and *AtCBF3* (Novillo *et al*., 2007). In many (but not in all) plant species signal diversification is also manifest as a fourth member of the CBF family. This CBF4 differs in its kinetic behavior. Its induction is slower than for the other CBFs, but persists longer, for instance in leaves and buds of grapevine (Xiao *et al*., 2008). The function of CBF4 is, therefore, linked with cold acclimation. Indeed, constitutive overexpression of *VvCBF4* produced grapevines with superior freezing tolerance (Tillett *et al*., 2012).

In our previous work (Wang *et al*., 2019), we found a correlation between the expression of grapevine CBF4 and the acclimation to cold stress. To test, whether this correlation derives from a causal relationship, we generated transgenic lines in the cellular model tobacco BY-2 that overexpressed CBF4 from the grapevine variety Pinot Noir in fusion with GFP. The CBF4-GFP resides in the cytoplasm, but shuttles to the karyoplasm in response to cold stress accompanied by improved cellular resilience. The inhibition of nuclear import by GTP-γ-S, exerts a strong effect on the response of CBF and COR transcripts to freezing. In this study, chilling and freezing stress differ with respect to the induction of transcripts for Cold-Box Factors and Cold Responsive (COR) genes, reflected by differences in early signalling. The swift and strong induction of CBF4 under freezing is a hallmark of tolerance, and that the responses to chilling are uncoupled from those to freezing. We infer from these data that chilling and freezing stress trigger different signal chains culminating in differential nuclear import of CBF4 leading to adaptive responses that depend on the type of cold stress.

## Materials and methods

### Cloning and stable transformation of *VvCBF4* into tobacco BY-2

Leaves of grapevine (*Vitis vinifera* L. cv. Pinot Noir, clone PN40024, the genotype used for the grapevine reference genome (Jaillon *et al*., 2007), were collected after exposure to 4 °C for 6 h. Total RNA was extracted using a commercial RNA isolation kit (Sigma Aldrich, Deisenhofen, Germany), and reversely transcribed into cDNA from 1 μg of mRNA template using the M-MuLV cDNA Synthesis Kit (New England Biolabs, Frankfurt am Main, Germany) according to the protocol of the manufacturer. Full-length *VvCBF4* (GenBank KF582403.1) was amplified with the modified oligonucleotide primers listed in **Supplementary Table S1** for subsequent integration into the GATEWAY pDONR/Zeo vector system (Invitrogen) using initial denaturation for 7 min at 94°C, and 30 cycles of denaturation for 1 min at 94°C, annealing for 1 min at 60°C, and elongation for 2 min at 72°C, with a terminal elongation step for 4 min at 72°C employing a proofreading Taq polymerase (Q5 High-Fidelity DNA polymerase, New England Biolabs, Germany). After verification of the amplicon sequence, the insert was then transferred into the binary vector pK7WGF2.0 making use of the recombinase function of the Gateway system (BP clonase II mix, Invitrogen Corporation, Paisley, UK). This construct allows for the expression of *VvCBF4* as fusion with green fluorescent protein (GFP) at the C-terminus is driven by the CaMV 35S promoter. The fusion construct was introduced into suspension cultured tobacco *(Nicotiana tabacum)* BY-2 cells using a strategy based on *Agrobacterium tumefaciens* (strain LBA4404) according to Klotz and Nick (2012). After co-cultivation for 3 d, droplets of the cell suspension were transferred on selective plates with 100 mg/L cefotaxime and 100 mg/L kanamycin. Upon four weeks of selection in the dark at 25°C, cells from a single growing callus were transferred into liquid BY-2 medium (see below) and then replated and re-picked on selective plates for three times before being used for the experiments.

### Cell culture and cold treatment

Tobacco *(Nicotiana tabacum* L. cv. Bright Yellow 2, BY-2) cells were cultivated as suspension in modified liquid Murashige and Skoog (MS) medium and sub-cultured weekly as described in (Schneider *et al*., 2015). To administer freezing (0°C) the entire Erlenmeyer flask with the cells was placed at day 5 after subcultivation into an ice-water mixture on an orbital shaker (KS260, IKA Labortechnik, Germany) at 150 rpm in the dark. To administer a chilling treatment (4°C), cells were kept in a cold room, again on an orbital shaker. If not stated otherwise, the cells remained exposed to the cold treatment for 24 h.

### Monitoring the cellular response to cold stress

To investigate the effect of *VvCBF4* on cold response, cell mortality of a time-course was assessed via Evans blue assay. Cells under freezing and chilling treatment were collected every 24h to continuously monitor cell mortality. 200μl cells were transferred into a mesh-like fixed with Polyamide fabric membrane (PA-41/31, Franz Eckert GmbH, Germany) and put on the tissue paper to remove the medium. Then the chambers were incubated in 2.5% (w/v) Evans Blue for 5 min followed by 5 times washing with distilled water. The dead cells were counted via microscopy by bright-field illumination. Data from each treatment at one time point were based on at least 4 independent experiments with more than 500 cells.

Extracellular alkalinization as a readout of calcium channel was performed. The changes in extracellular pH were measured by a pH meter (pH 12, Schott Handylab) with a pH electrode (LoT 403-M8-S7/120, Mettler Toledo). The cell suspension (4 mL) was pre-equilibrated on a shaker at 25°C in the dark for 1 h, before administering cold stress. Besides, the intracellular distribution of calcium was measured by the fluorescent dye chloro-tetracycline. The cells were sampled after the respective treatment and then transferred into a mesh-like device for removing the medium. The filtered cells were fixed in 2.5% glutaraldehyde in 200 mM sodium phosphate buffer (pH 7.4) for 15 min. The fixative was washed out in three rounds (each 5 min) with staining buffer (50 mM Tris-HCl, pH 7.45), and excess liquid was carefully drained out by filter paper, before staining for 5 min with 100 μM chlorotetracycline. Unbound dye was washed out twice for 2 min, and cells were directly analyzed by spinning-disc confocal microscopy. The green fluorescence was recorded by spinning disc confocal microscopy upon excitation with the 488-nm line of an Ar-Kr laser (Zeiss) and collecting the green emission. For evaluation, mean fluorescence intensity was quantified using ImageJ. Concerning, the GFP emission of the VvCBF4ox line, Data was calculated by the fluorescence intensity of each time point minus the mean of the initial fluorescence intensity from the untreated OE-CBF4 line.

### Inhibition of transcription and translation

To test for the role of transcription in the regulation of CBF4, the inhibitor Actinomycin D (AMD, Sigma-Aldrich, Deisenhofen, Germany) was used, which intercalates into the DNA and, thus, prevents transcriptional elongation by polymerase I (for review see (Bensaude, 2011)). To address the role of translation, we used the protein synthesis inhibitor cycloheximide (CHX, Sigma-Aldrich, Deisenhofen, Germany). CHX inhibits eEF2-mediated translocation in eukaryotic ribosomes (Schneider-Poetsch *et al*., 2010). Both, VvCBF4 over-expressor, and non-transformed BY-2 wildtype cells were pre-treated for 2 h with either 100 μg/ml CHX or 50 μg/ml AMD under shaking at 150 rpm in the dark, before transfer to 0°C and 4°C for subsequent 24 h. As solvent control for AMD, cells were treated with 2.5% ethanol. Cells were sampled directly after the pre-treatment, and cells treated with cold only without inhibitor pre-treatment were included as additional controls.

### Pharmacological treatments

To address the role of the NADPH oxidase Respiratory burst oxidase Homologue (RboH) in cold signaling, cells were pre-treated with 10 μM of Diphenyleneiodonium (DPI) specifically inhibiting the generation of superoxide (Eggenberger *et al*., 2017). The MAPK cascade was assessed by pre-treatment with 100 μM of the MAP kinase kinase blocker PD98059 (Zhang *et al*., 2006). The cold-induced transcription factor cascade is under control of the regulator Inducer of CBF expression 1 (ICE1), a protein that is rapidly generated and broken down under normal temperature. In contrast, this protein is accumulating in the cold, because its proteolysis in the proteasome is blocked. To address this, we used 100 μM of MG132, a specific inhibitor of the proteasome (Jang *et al*., 2005). All three inhibitors were purchased from (Sigma-Aldrich, Deisenhofen, Germany), and dissolved from stock solutions in DMSO. They were administered 30 min prior to cold-treatment, and the cells were incubated at 27°C on an orbital shaker (KS260, IKA Labortechnik, Germany) at 150 rpm in the dark. After pretreatment, cells were transferred to 0°C or 4°C, respectively, for 24 h on the shaker in the dark at 150 rpm. For the solvent control, cells were treated with 1% (v/v) DMSO, corresponding to the solvent concentration in the inhibitor treatments. Additionally, cells without pharmacological treatment under 0°C and 4°C for 24 h were added as reference.

### Inhibition of nuclear import by GTP-γ-S

To address the role of nuclear import, the non-hydrolysable GTP analogue GTP-γ-S was used. This compound blocks the GTPase function of Ran also in plants (Merkle *et al*., 1996) and, thus, should block a potential nuclear import of VvCBF4ox. Cells were pre-treated for 30 min with 500 μM GTP-γ-S dissolved in distilled water at 25°C, prior to cold stress (0°C or 4°C) for 24 h on the shaker in the dark at 150 rpm. Control cells were treated in the same manner, but omitting GTP-γ-S.

### Inhibition of jasmonate biosynthesis by phenidone

Since Jasmonate-ZIM domain (JAZ) proteins, the repressors of jasmonate signalling, were reported to interact with ICE1 and suppress its transcriptional activity (Hu *et al*., 2013), we used 1-phenylpyrazolidinone (phenidone) to inhibit jasmonate biosynthesis. Phenidone blocks the conversion of α-linolenic acid into the jasmonate precursor 13-HPOT (Bruinsma *et al*., 2010a). Cells were pre-treated with 2mM phenidone at room temperature for 30 minutes on the shaker in the dark at 150 rpm and subjected to cold stress (either 0°C or 4°C) for the subsequent 24 h. As solvent control, cells were treated with same volume ethanol, corresponding to the solvent concentration in the inhibitor treatments. Additionally, cells without pharmacological treatment under 0°C and 4°C for 24 h were added as mock controls.

### Quantification of *CBF4* transcripts in grapevine leaves under cold stress

To get insight into the functional context of CBF4 induction, we used a series of grapevine species differing with respect to their tolerance. These included the cold-tolerant *V. amurensis* (KIT-voucher 6540), the cold-sensitive *V. coignetiae* (KIT-voucher 6542), and *Vitis vinifera* L. cv. Pinot Noir (KIT-voucher 7474, corresponding to clone PN40024 that had been used for the reference genome (Jaillon *et al*., 2007)) from the germplasm collection established in the Botanical Garden of the Karlsruhe Institute of Technology. Plantlets were propagated clonally from wood cuttings and were used for this experiment in the age of 10 weeks. Entire plants were transferred either to a chilling treatment (in a cold room at 4°C) or subjected to a freezer (−18°C) for specific time intervals (0.1, 3, 6, 12 and 24 h). Then, the first fully expanded leaves (plastochrones 5-7) were excised, immediately frozen in liquid nitrogen, and the samples stored at −80°C for further investigation.

### Quantitative real-time PCR (RT-qPCR) analysis

Total RNA was extracted from BY-2 suspension cells using the innuPREP Plant RNA Kit (Analytik Jena, Jena, Germany). For extraction from leaves, the Spectrum™ Plant Total RNA Kit (Sigma, Germany) was used. Complementary DNA was synthesised with the M-MuLV Reverse Transcriptase (New England Biolabs, Frankfurt, Germany) and used as a template for quantitative Real-Time PCR (qRT-PCR) as described in (Wang *et al*., 2018). The details including the sequence of the oligonucleotide primers are listed in **Supplementary Table S1**. Steady-state transcript levels were calculated according to (Livak and Schmittgen, 2001) and normalised against L25 as reference gene. Data represent mean and standard errors from three independent experiments, each in technical triplicates.

## Results

### *VvCBF4* is imported into the nucleus in response to cold stress

To obtain insight into the function of VvCBF4, we generated a C-terminal fusion with GFP and expressed this fusion construct in tobacco BY-2 cells under control of the constitutive CaMV 35S promoter. Upon transient expression following biolistic transformation and inspection by spinning-disc confocal microscopy, we observed a punctate signal in the perinuclear cytoplasm and the transvacuolar strands that emanate from the nucleus in these vacuolated cells (**Fig. 1A, B**). In contrast, the karyoplasm was void of any signal. Since *VvCBF4* (as well as the other members of the CBF family) harbored a *bona-fide* nuclear localisation signature (**Fig. S1**), we would have expected an intranuclear GFP signal. To address this further, we generated a stable transgenic line expressing the VvCBF4-GFP fusion. When these cells were investigated at normal temperature (25°C), we observed again a cytoplasmic signal (**Fig. 1C**). When we zoomed into the GFP-channel and compared it with the overlay with the image obtained by Differential Interference Contrast (**Fig. 1D**), the GFP signal was located clearly outside of the nuclear envelope, while no signal was detectable in the karyoplasm, confirming the results from the transient expression. We wondered, whether the subcellular localisation might be conditional, then conducted an experiment, where the cells were subjected to severe cold stress (0°C for 4 h). This resulted in a generally reduced intensity of the signal (**Fig. 1E**), but now, the close-ups revealed that the protein was found in speckles localised inside of the nuclear envelope (**Fig. 1F**). Thus, in the absence of cold stress, the GFP fusion of VvCBF4 is cytoplasmic, but seems to be imported into the nucleus, when the cells experience cold stress.

**Figure 1.**
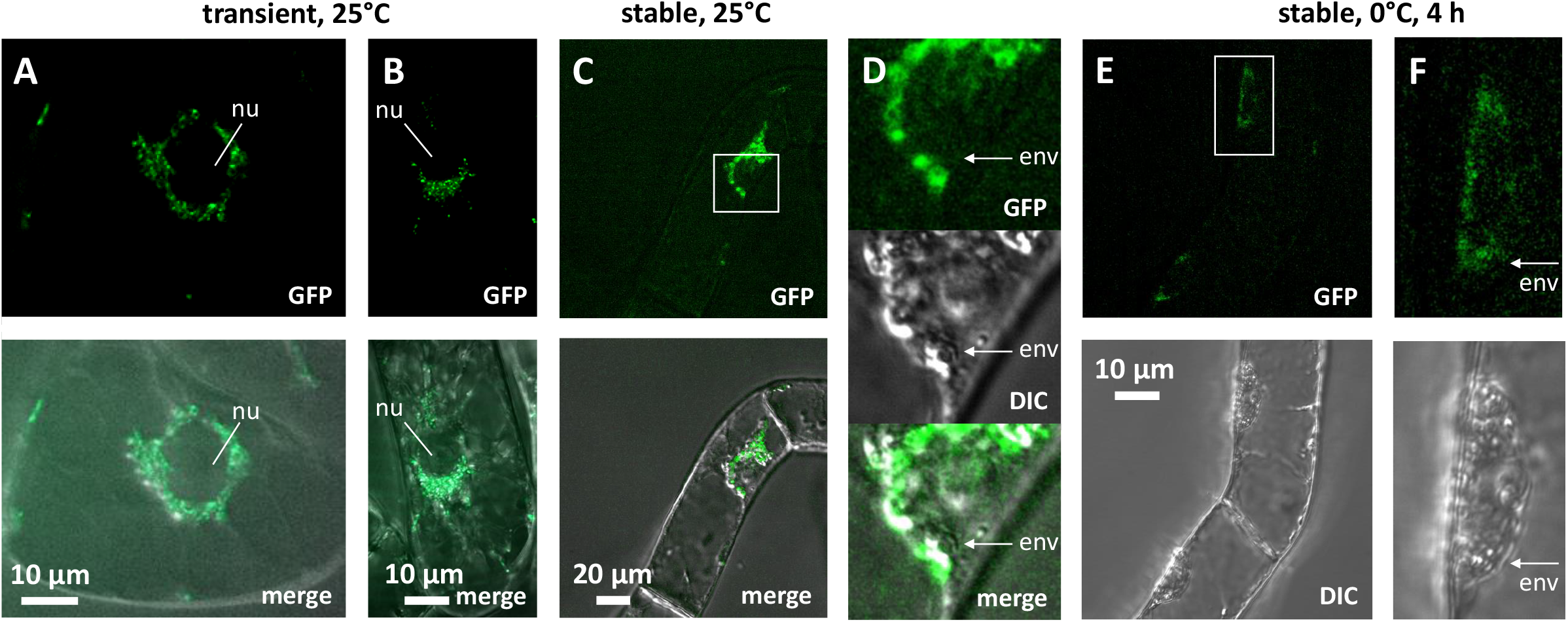
Subcellular localisation of VvCBF4 in fusion with GFP, assessed in tobacco BY-2 cells as heterologous host. **A, B** transient transformation with a high-copy vector, cells incubated at 25°C. For two representative cells the GFP signals and the merge of the GFP signal with the differential interference contrast (DIC) are shown. Note that the GFP signal is localised outside of the nucleus (nu). **C, D** stable transformation with a binary vector, cells incubated at 25°C. **D** shows a zoom-in of the region marked by a white square in **C**. White arrows indicate the position of the nuclear envelope (env) separating karyoplasm and cytoplasm to show that the GFP signal is found outside of the nucleus. **E, F** stable transformation with a binary vector, cells incubated at 0°C for 4 h. **F** shows a zoom-in of the region marked by a white square in **E**. White arrows indicate the position of the nuclear envelope (env) separating karyoplasm and cytoplasm to show that this time the GFP signal is found inside of the nucleus.

### *VvCBF4* mitigates mortality inflicted by cold stress

To address the role VvCBF4 for cold tolerance, we followed cell mortality under continuous cold stress either in the BY-2 cell line stably expressing *VvCBF4,* or in the non-transformed wildtype (WT) (**Fig. 2**). When the WT was kept at 0°C, mortality increased steeply from 24 h after induction increasing 3 to 4-fold over the resting level within 72 h. For VvCBF4 overexpressor, this cold induced mortality was clearly mitigated by around 40% at 72 h, while the mortality of unchallenged cells was comparably low for both lines. In the next step, we followed mortality under chilling stress (4°C). Here, a significant increase of mortality was observed much later (from 72 h of stress treatment) and to a lower amplitude (less than 2-fold over the resting level at 72 h of chilling. Again, VvCBF4 overexpressor seemed to be sturdier, but the difference was less pronounced (by around 20% at 72 h) and not reaching significance. Thus, expression of VvCBF4 mitigates mortality inflicted by cold stress.

**Figure 2.**
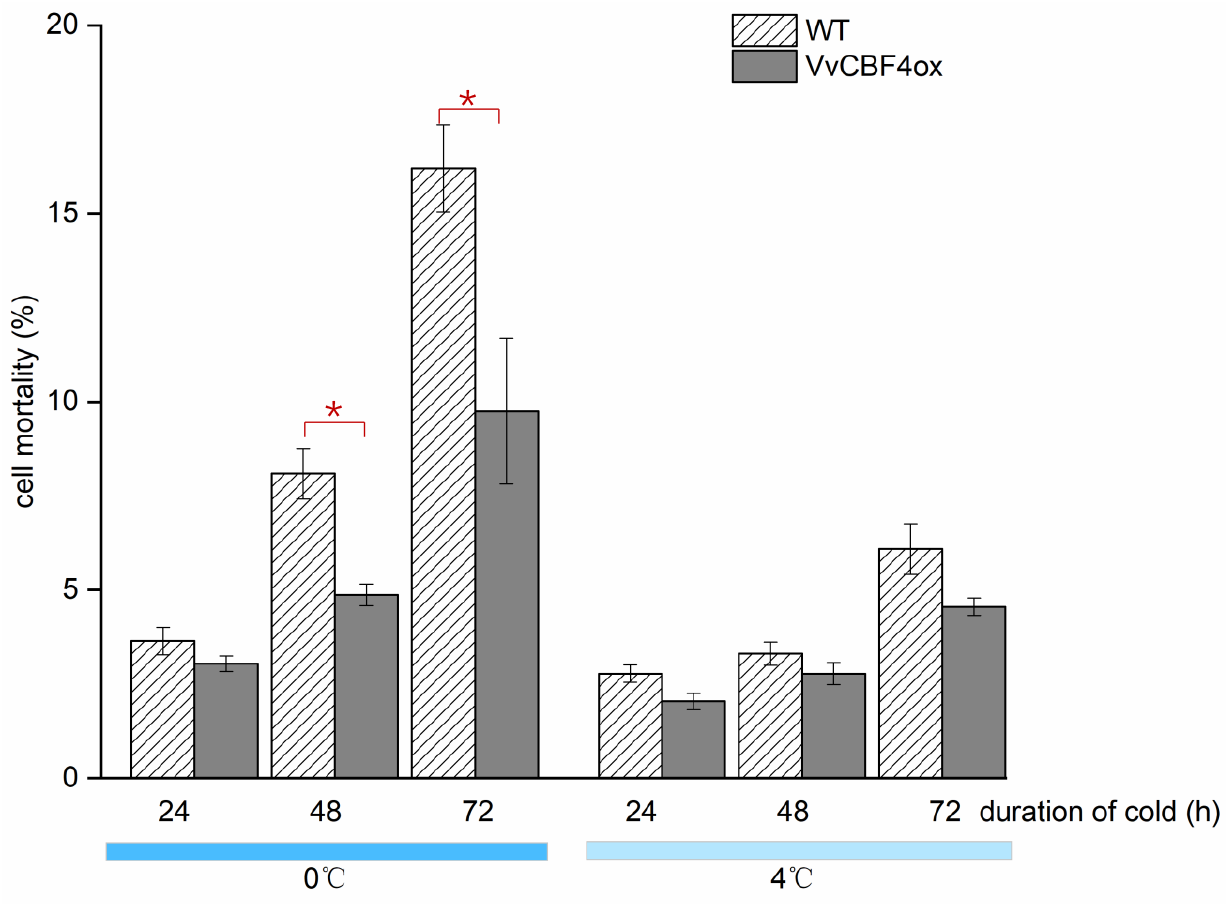
Cell mortality in response to cold treatment (4 °C and 0 °C). Data are shown as means±SE from three independent experiments with 1500 cells each. Asterisk represents a statistically significant difference with Fisher’s LSD test (P <0.05).

### *VvCBF4* and its endogenous homologue are regulated differently in tobacco BY-2

To understand, why the transfected VvCBF4 mitigated cold-inflicted mortality in the tobacco BY-2 recipient, we followed steady-state transcript levels of *VvCBF4* along with transcripts of the tobacco CBF4 paralogue, *Avr9/Cf9,* over time in response to both chilling (4°C) and severe cold stress (0°C). The resting level of *Avr9/Cf9* was identical in WT cells and in VvCBF4 overexpressor (**Fig. 3A**), indicating that the overexpression left the expression of the endogenous CBF4 paralogue unaltered. The resting level of the transfected *VvCBF4* was around 6.5 times higher as compared to the endogenous *Avr9/Cf9*, which is to be expected, since *VvCBF4* is driven by the CaMV 35S promoter. Interestingly, the steady-state level of *VvCBF4* transcripts increased significantly, when the overexpressor cells were exposed to cold stress (**Fig. 3B**). This induction was more pronounced for chilling as compared to severe cold stress. For chilling, the *VvCBF4* transcripts increased by 3.5-fold of the resting level (corresponding to around 25 times the resting level of *Avr9/Cf9),* but only 2-fold for severe cold stress (around 13 times the resting level of *Avr9/Cf9*).

**Figure 3.**
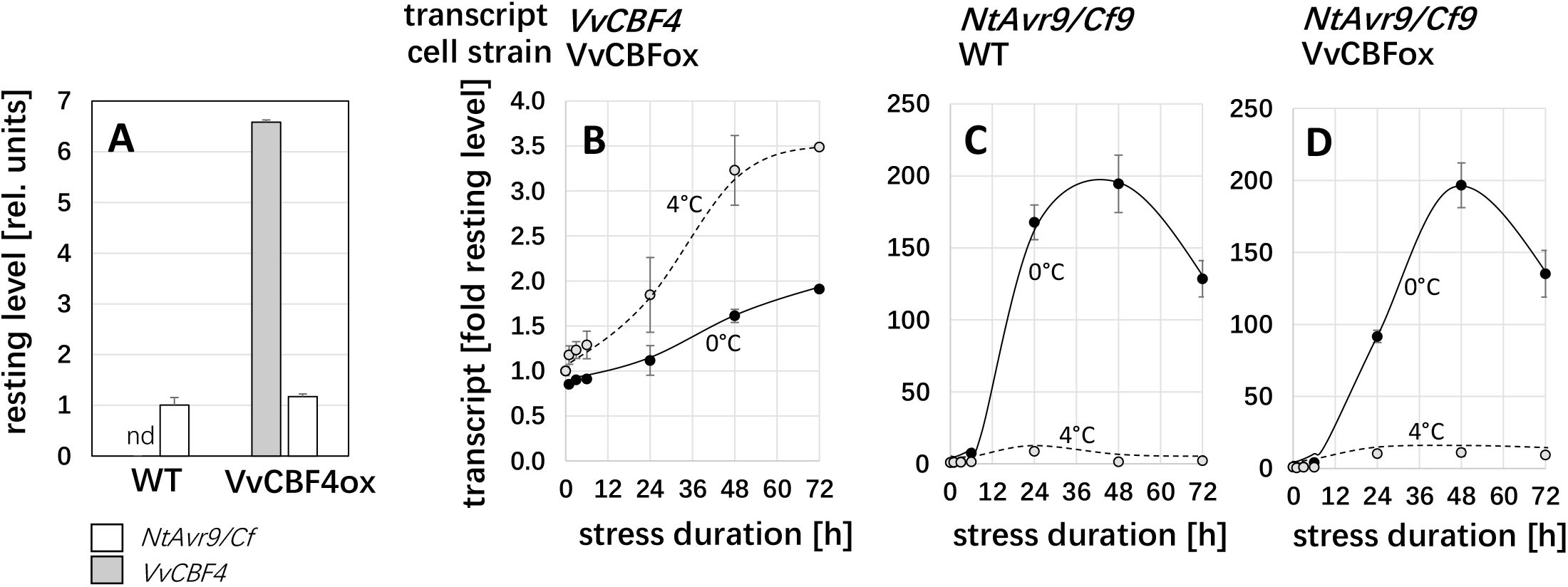
Regulation of transcripts for *VvCBF4* and its tobacco homologue *NtAvr9/Cf9* in response to chilling (4°C) and freezing (0°C) depending on cell strain. A resting levels for the transcripts of the two CBF4 homologues in the absence of cold stress in non-transformed BY-2 cells (WT) versus cells constitutively overexpressing VvCBF4 in fusion with GFP (VvCBF4ox). nd non detectable. **A-C** Time course for steady-state transcript levels for *VvCBF4* **(B)** and native *NtAvr9/Cf9* **(C, D)** during chilling and freezing in the background of the non-transformed WT **(C)**, or cells overexpressing VvCBF4 **(B, D)**. Data are expressed relative to the resting levels (given in **A**) and represent mean±SE from three independent experimental series, each in technical triplicates.

In the next step, we investigated the regulatory pattern for the tobacco paralogue of *CBF4*, *Avr9/Cf9,* under the same conditions. Irrespective of the absence (**Fig. 3C**) or the presence (**Fig. 3D**) of the transgene, there was a strong induction of *Avr9/Cf9* transcripts peaking at around 200-fold of the resting level at 48 h after the onset of severe cold stress (0°C). For chilling stress, only a relatively minute induction of less than 10-fold over the resting level resulted. Thus, the temperature dependency of the foreign *VvCBF4* (stronger induction at 4°C compared to 0°C) was inverted to the pattern seen for the endogenous *Avr9/Cf9* (weaker induction at 4°C compared to 0°C). The regulation of the endogenous *Avr9/Cf9* was not affected by the overexpression of *VvCBF4*.

### *VvCBF4* modulates expression of specific CBFs and CORs depending on temperature

The transcripts of the foreign *VvCBF4* were upregulated under cold stress (**Fig. 3B**), which was accompanied by a reduced mortality (**Fig. 2**). As to get insight into the potential functions of *VvCBF4* during the response to cold stress, we followed the expression of the endogenous CBF factors *DREB1* and *DREB3*and the three COR transcripts *NtERD10a, c*and *d*(**Fig. 4**).

**Figure 4.**
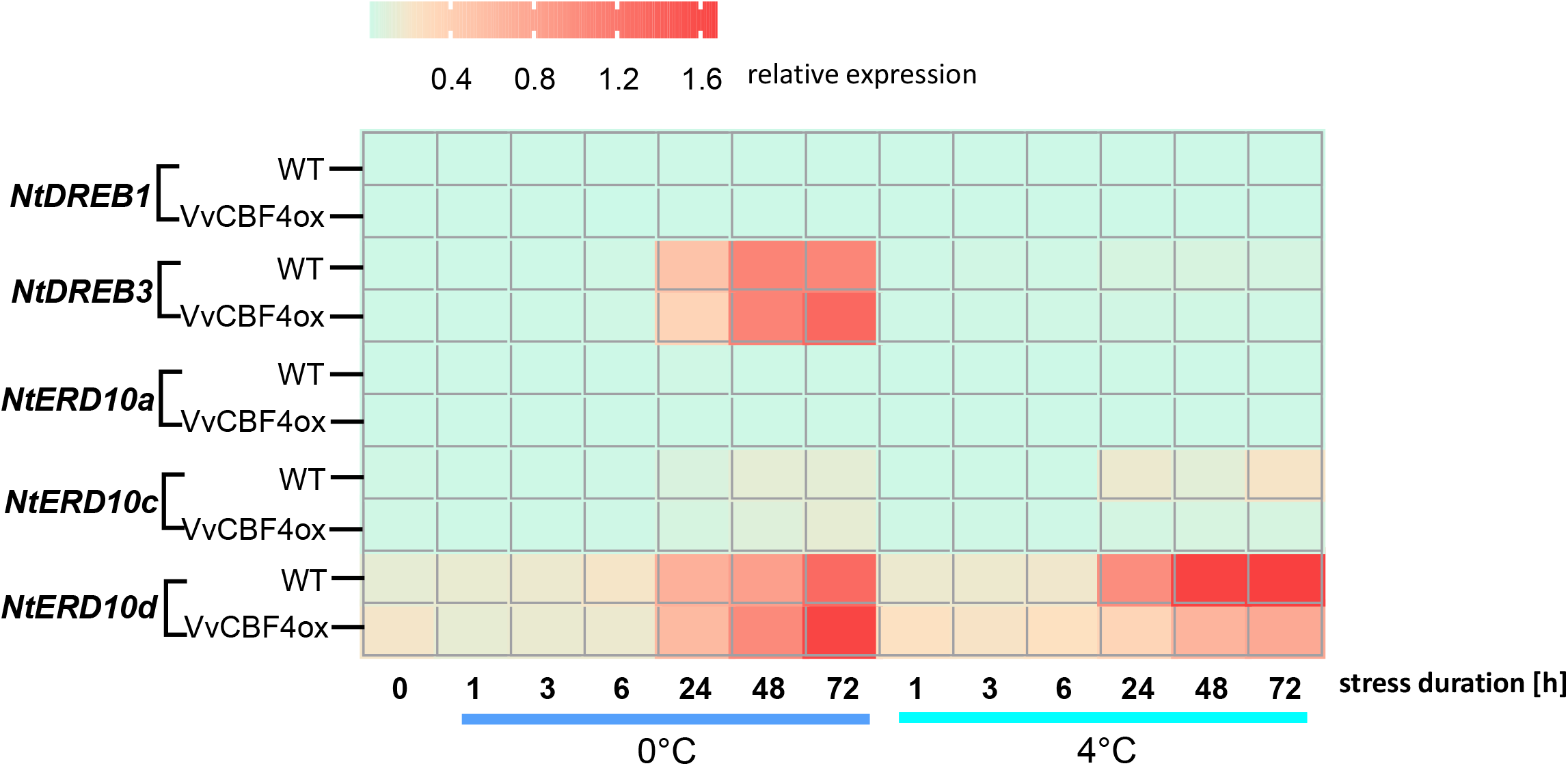
Time course of relative expression of endogenous CBF genes and COR genes in response to chilling (4°C) and freezing (0°C). Colour code shows steady-state levels for two endogenous CBF *(NtDREB1* and *NtDREB3)* and three COR transcripts *(NtERD10a, NtERD10c* and *NtERD10d)* relative to L25 based on 2^-ΔCt^. Data represent means from three independent experimental series, each in technical triplicates.

The differences between *DREB1* and *DREB3* were quite remarkable (**Fig. 4**). While *DREB1* transcripts did not display any noteworthy changes, neither in the WT, nor in the VvCBF4 overexpressor, neither for freezing, nor for chilling, there was a conspicuous response of *DREB3* transcripts. These increased strongly from 24 hours after the onset of freezing stress, while there was no induction in case of chilling stress. This accumulation of *DREB3* transcripts reached a plateau at 118-fold of the resting level from 48 h in case of the WT, while in the VvCBF4 overexpressor it continued to grow further to reach 150-fold after 72 h of freezing stress.

Since cold tolerance in plants largely relies on the induction of COR proteins that are activated by upstream transcription factors, such as the CBFs, we examined the dehydrin transcripts for *NtERD10a* (GenBank AB049335) coding for a late embryogenesis abundant (LEA) protein, and *NtERD10c* (GenBank AB049337) and *NtERD10d* (GenBank AB049338), representing splice variants of the same gene, encoding a ECPP44-like phosphoprotein. While there was no significant response of *NtERD10a* for any of the cell lines or stress conditions, transcripts for *NtERD10c* and *d* did respond clearly. Transcripts for *NtERD10c* were induced after 24 h of chilling in the WT, while in the VvCBF4 overexpressor no induction was seen. In case of freezing, the induction in the WT was slower and of lower amplitude (after 72 h, an induction by 6-fold was observed). In the VvCBF4 overexpressor, this induction was even further delayed, although the induction at 72 h was comparable to that of the WT (around 8-fold of the resting level). The most salient response was observed for transcripts of *NtERD10d* (the splicing alternative of *NtERD10c*). These were strongly induced from 24 h of cold stress in both cell lines under both, freezing and chilling stress. However, the amplitude of this response was inverted in the two cell lines: In the non-transformed WT, an induction of 8.7-fold was reached after 72 h of freezing stress, while for the VvCBF4 overexpressor, this induction was even stronger (10-fold at 72 h of freezing stress). Under chilling stress, the induction was more pronounced in the WT (14-fold at 72 h), while it was now much weaker (only 4-fold at 72 h) in the VvCBF4 overexpressor. This inverted temperature dependency between the lines is matching with the inverted pattern seen for the foreign *VvCBF4* transcript (**Fig. 3B**) as compared to the endogenous CBF4 orthologue *Avr9/Cf9* (**Fig. 3C**). Taken together, the overexpression of VvCBF4 amplifies the accumulation of *DREB3* transcripts in response to freezing stress accompanied by elevated accumulation of the *ERD10d.* In the WT, this splice variant is preferentially induced by chilling (not by freezing), correlating with the pattern of the endogenous CBF4 orthologue, *Avr9/Cf9.* In other words, the introduced *VvCBF4*imposes the temperature sensitivity pattern of its donor, *Vitis vinifera,* on the *COR* transcript of the recipient, *N. tabacum*.

### Nuclear import modulates the response of CBFs and COR to cold stress

As we had observed that *VvCBF4* is translocated from the cytoplasm into the nucleus in response to cold stress (**Fig. 1**), and since ectopic overexpression of *VvCBF4* altered the response of endogenous CBFs and COR genes to cold stress (**Fig. 4**), we questioned, whether translocation of *VvCBF4* into nucleus is required for this modulation of gene expression. To address this, we tested the effect of GTP-γ-S, which irreversibly blocks the GTPase function of Ran, and had been shown to block nuclear import also in plant cells (Merkle *et al*., 1996). In the absence of cold stress, with exception of a slight inhibition of *DREB1* transcripts, none of the tested transcripts displayed any change (**Fig. 5**). However, under cold stress, the expression of the tobacco CBF4 homologue *Avr9/Cf9* was modulated – the induction under freezing stress was reduced to ¼ after pre-treatment with GTP-γ-S indicating that nuclear import is essential for this induction by freezing stress. The relatively low induction of *Avr9/Cf9* transcript by chilling stress was not altered. In contrast, transcripts for *NtDREB1* that are downregulated around 5-fold by freezing stress, remain unaltered in presence of GTP-γ-S. Again, chilling caused only comparatively small effects that were reduced further by the inhibitor. The transcripts for the highly responsive *NtDREB3* were reduced from a 30-fold to a 16-fold induction by GTP-γ-S under freezing stress (under chilling stress, this transcript did not change significantly, neither for absence nor presence of the inhibitor). Thus, *NtDREB3* behaved parallel to *Avr9/Cf9.* For the COR gene *NtERD10a,* the inhibitor did not cause significant changes in the response to cold stress. However, *NtERD10c* and *NtERD10d* were altered clearly and with the same pattern by GTP-γ-S. Here, the induction by freezing stress was not changed, but the induction by chilling stress was boosted by around twofold. Thus, nuclear import supports the induction of the CBF4 orthologue *Avr9/Cf9* and the CBF *NtDREB3,* while it acts negatively upon the induction of the COR transcripts *NtERD10c* and *d*

**Figure 5.**
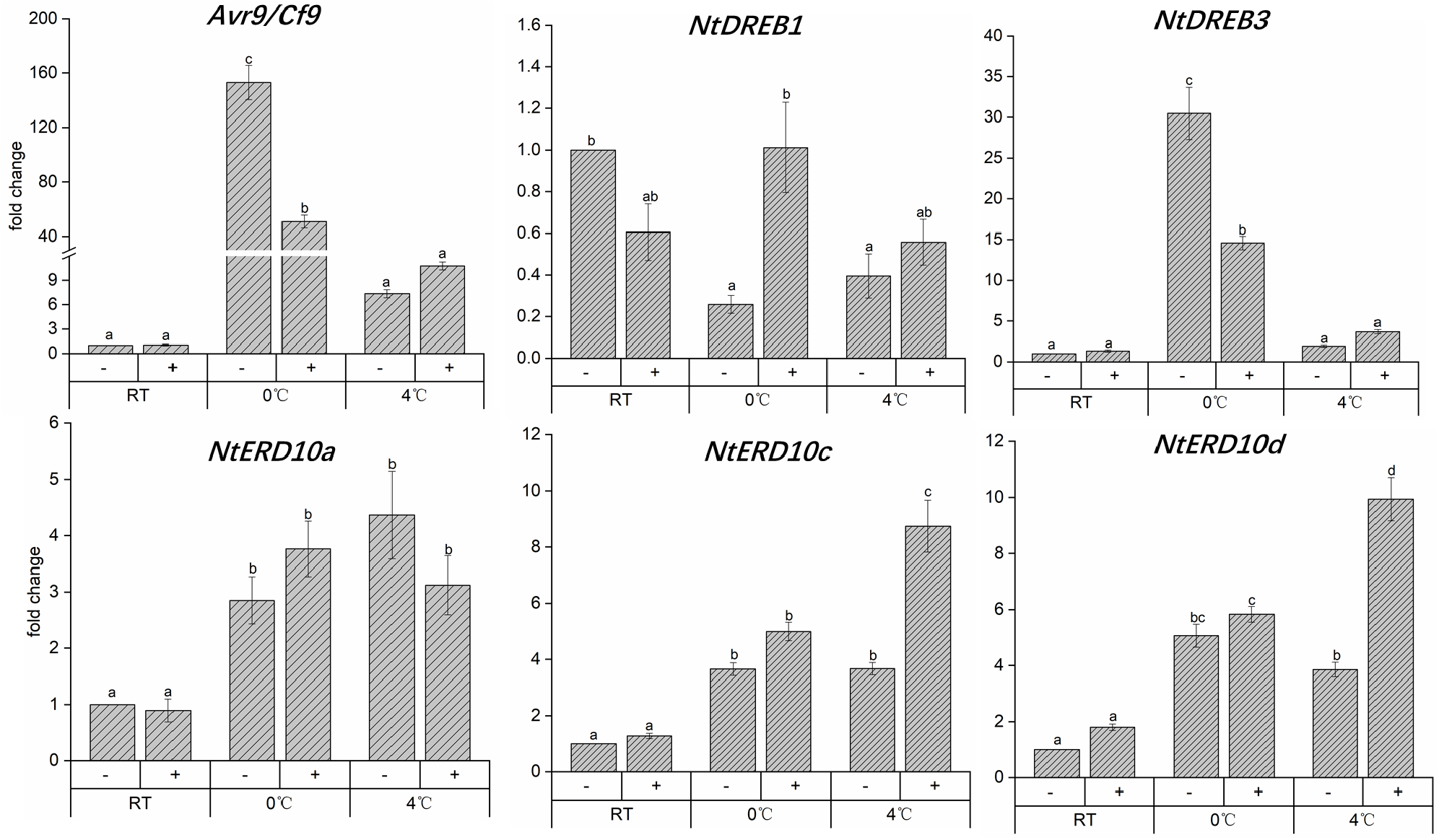
Effect of the nuclear-import inhibitor GTP-γ-S on the cold responses of CBF and COR transcripts in the VvCBF4 overexpressor line, measured by quantitative real-time PCR analysis for endogenous CBF and COR genes in response to chilling (4°C) and freezing (0°C). Cells were pre-treated with 500 μM GTP-γ-S for 30 minutes at 25°C prior to cold stress (0°C or 4°C) for additional 24 h and then examined for gene expression. Controls (−) were treated in the same way, but omitting GTP-γ-S, while + represents the actual test, where cells were treated with GTP-γ-S. Data represent means ± SE from three independent experiments conducted in technical triplicates and are normalised to the untreated control. Different letters indicate the statistically significant difference based on a LSD test (P < 0.01).

### The COR transcript ERD10d is negatively regulated by a protein synthetised *de-novo*

To understand the mechanism behind the differential effect of chilling and freezing stress upon the induction of *ERD10d* transcripts (that derive by differential splicing from the same gene as *ERD10c),* we probed for the effect of Actinomycin D (AMD) as inhibitor of transcription, and Cycloheximide (CHX) as inhibitor of translation. To find out, whether the treatment used in this experiment was sufficient, we first tested their effects on the transcripts of the foreign *VvCBF4* that are driven by the constitutive CaMV 35S promoter (**Fig. 6A**). Indeed, inhibition of transcription by AMD significantly reduced *VvCBF4* transcripts to a residual level of around 35% of the resting level, irrespective of temperature. In contrast, inhibition of translation by CHX did not modulate the strong induction of *VvCBF4* transcripts (around 3-fold) in response to chilling stress, nor did it alter the weak induction of these transcripts in response to freezing stress. The pattern for *ERD10d* transcripts (**Fig. 6B**) differed fundamentally. While AMD eliminated the induction of these transcripts irrespectively of temperature, there was a clear difference with respect to the effect of CHX. In the absence of cold stress, *ERD10d* transcripts were induced 3-fold by CHX. Likewise, the around two-fold induction of *ERD10d* transcripts in response to chilling was further accentuated by CHX to around 4-fold of the resting level. In contrast, the strong induction of *ERD10d* transcripts in response to freezing remained mostly unchanged after pre-treatment with CHX. Thus, while transcription is necessary for the induction of *ERD10d,* there seems to be a negative regulator (both in the absence of cold stress and under chilling stress) that is synthetised as protein *de-novo.* However, this negative regulator is not needed for the response of this transcript to freezing stress.

**Figure 6.**
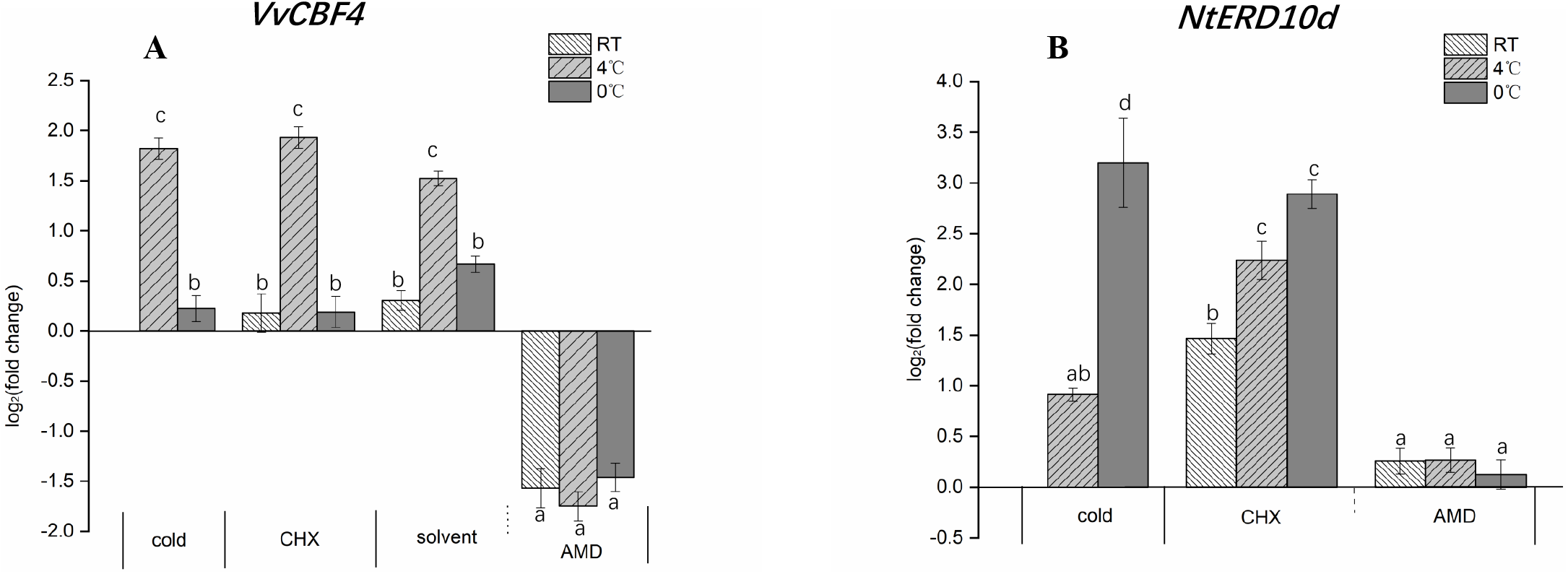
Role of transcription for expression of *VvCBF4* and *NtERD10d.* Steady-state transcript levels for the foreign *VvCBF4* (**A**) and the endogenous COR transcript *NtERD10d* (**B**) were measured after 24h incubation of VvCBF4 overexpressor cells and none-transformed BY-2 either at 25°C (RT), or chilling (4°C), or freezing (0°C) stress after pre-treatment for 2 h with either the transcription inhibitor Actinomycin D (AMD, 50 μg/ml) or the translation inhibitor Cycloheximide (CHX, 100 μg/ml). The solvent control of AMD was 2.5% EtOH, while CHX was diluted in an aequous stock. Data are normalised to the expression levels under room temperature without any of the inhibitors and represent means±SE of three independent experiments in technical triplicate. Statistically significant difference are represented as different letters with Fisher’s LSD test (*P* <0.01).

### Different CBF4-dependent signaling in chilling and freezing stress

To get insight into the signalling underlying the regulation of cold-related transcripts, we determined transcripts for *VvCBF4* (in VvCBF4 overexpressor) and *NtERD10d*(in both VvCBF4 overexpressor and non-transformed WT cells) after 24 h of either freezing (0°C) or chilling (4°C) stress pre-treated with different inhibitors (**Fig. 7**). The two stress treatments alone induced a different response of the two transcripts - *VvCBF4* was induced by chilling, but not by freezing stress, while *NtERD10dwas* more induced by freezing stress and less by chilling stress (**Fig. 7**, mock). While the response of *VvCBF4* did not show obvious changes in presence of the solvent (1% DMSO), that of NtERD10d was clearly modulated. The induction by freezing stress was partially silenced by this solvent in the VvCBF4ox line, while it was significantly enhanced in the non-transformed WT. For chilling stress, DMSO clearly enhanced the induction of *NtERD10d* in both cell lines. Thus, DMSO, which is not only a solvent, but also a membrane rigidifier, had a clear effect that was depending on the type of cold stress and the overexpression of *VvCBF4.* Pre-treatment with Diphenyliodonium (DPI), an inhibitor of the NADPH oxidase Respiratory burst oxidase Homologue, significantly quenched the induction of *VvCBF4* transcripts by chilling stress, while *NtERD10d* was not affected (neither in the overexpressor line or WT). Inhibition of MAPK signalling by PD98059 suppressed the chilling response of *VvCBF4* transcripts even stronger. Here, we observed a mild (in the overexpressor line) or strong (in the WT) induction of *NtERD10d* transcripts. Again, the responses to freezing stress were modulated much less. The most salient changes were seen with MG132, a specific inhibitor of the proteasome. Here, both, *VvCBF4*and *ERD10d,* were strongly induced under chilling stress. A comparable induction of *ERD10d* transcripts was seen for freezing stress, while *VvCBF4* transcripts did not exhibit this induction. Overall, the induction of *VvCBF4* transcript by chilling not only requires RboH and MAPK signalling but is also boosted by a factor that is swiftly degraded by the proteasome. This factor is not playing a role in freezing stress. This factor also boosts the accumulation of *ERD10d,* in contrast to *VvCBF4,* during freezing stress as well. These data show that the response of the COR transcript depends on the presence of the foreign *VvCBF4* gene, and that the signalling differs qualitatively between chilling and freezing stress. The most striking trait is the strong dependence of the transcripts on the proteasome because its inhibition by MG132 was strongly enhancing the expression of both cold-related transcripts under chilling stress, in case of *ERD10d* also under freezing stress.

**Figure 7.**
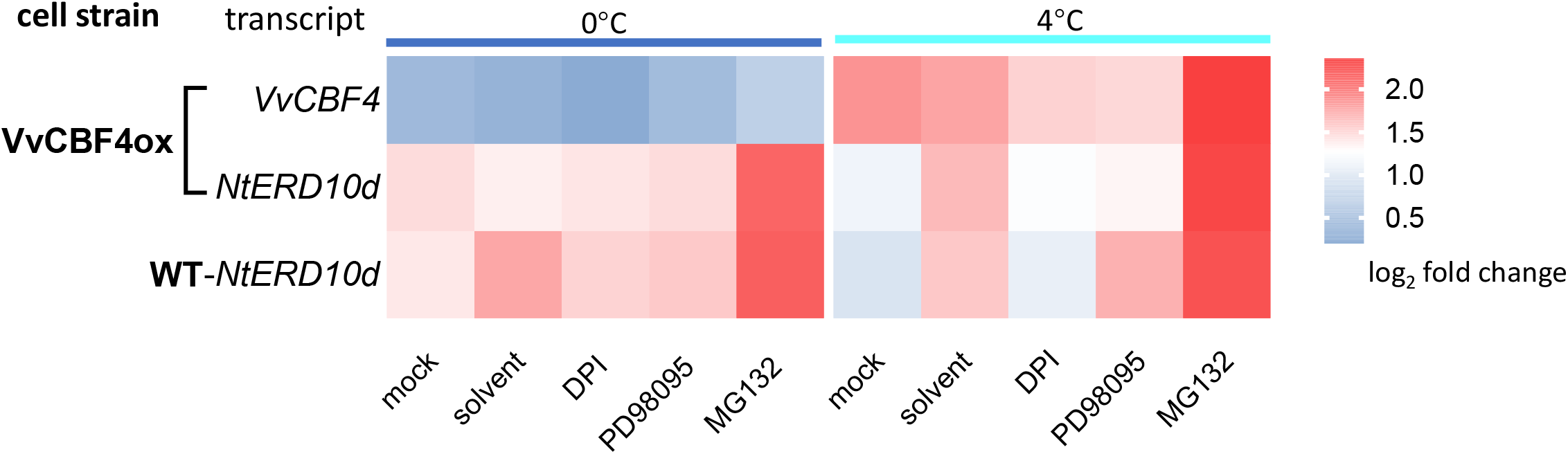
Response of transcripts for *VvCBF4* (in tobacco VvCBF4ox cells) and *NtERD10d* (in both, VvCBF4ox and non-transformed BY-2 cells, WT) to 24 h of freezing (0°C) or chilling (4°C) stress after pre-treatment for 30 min with either 10 μM Diphenyleneiodonium (DPI, inhibiting the NADPH oxidase Respiratory burst oxidase Homologue), or 100 μM PD98059 (inhibiting MAPK signalling), or 100 μM MG132 (inhibiting the proteasome). For the solvent control, cells were pre-treated with 1% (v/v) DMSO in the same volume in treatments. The mock control consisted in an experiment, where the cells were subjected to the respective cold stress for 24 h in the absence of any inhibitor. Data are normalised to the expression levels under room temperature without any of the inhibitors and represent means±SE of three independent experiments in technical triplicate. Statistically significant difference are represented as different letters with LSD test (*P* <0.01).

### The lipoxygenase inhibitor phenidone induces *VvCBF4* and *VvCBF4*-dependent *NtERD10d*

The strong effect of the proteasome inhibitor MG132 (**Fig. 7**), led us to the question, whether this effect might be related to the continuous turnover of ICE1. The transcriptional activity of this master switch was shown for *Arabidopsis thaliana* to be modulated by physical interaction with specific members of the JASMONATE ZIM DOMAIN family (Hu *et al*., 2013). These proteins are negative regulators of jasmonate signalling but are produced in response to jasmonate signalling. If this very specific hallmark of cold signalling would be conserved in tobacco cells, we should see specific changes in the cold response, when the production of jasmonates is disrupted. This can be achieved by phenidone, an inhibitor of lipoxygenase, thus, blocking the conversion of α-linolenic acid into the jasmonate precursor 13-HPOT (Bruinsma *et al*., 2010a; Bruinsma *et al*., 2010b). The solvent control, 1% ethanol exerted some effect as well, *VvCBF4* expression was induced by 2-fold at 25°C (**Fig. 8A**). Likewise, at 25°C *NtERDlūd* expression was induced by 4-fold in the VvCBF4 overexpressor, and even 13-fold in the WT (**Fig. 8B**). Under cold stress, this effect of the solvent was not noted with exception of a slight, but significant enhancement for the freezing response of *NtERD10d* transcripts. However, the effects of the solvent were minor as compared to the effect seen for phenidone, which already under room temperature stimulated the transcripts of *VvCBF4* by 4-fold (**Fig. 8A**). The transcripts of *ERD10d* were elevated as well to a similar degree (in the WT by around 4-fold, in the VvCBF4ox by around 6-fold) (**Fig. 8B**). Under chilling, phenidone did not yield conspicuous effects, for *VvCBF4,* transcript levels were even exactly the same as in the mock control (**Fig. 8A**). The situation differed drastically for freezing stress. Here, *VvCBF4* was induced 15-fold (**Fig. 8A**). For *NtERD10d*, the induction was even more pronounced, but only when *VvCBF4* was overexpressed. Under these conditions, *NtERD10d* was induced more than 50-fold (**Fig. 8B**), comparing to around 10-fold in the absence of phenidone. Interestingly, this induction was not seen in theWT (lacking the foreign VvCBF4 gene and, thus, the induction of VvCBF4 transcripts by phenidone (**Fig. 8A**). Taken together, we can show that jasmonates modulate the induction of *VvCBF4* and *ERD10d* transcripts under freezing stress. The fact that there is no such induction of *ERD10d* seen in WT, indicates that the endogenous CBF4 orthologue *Avr9/Cf9* cannot functionally replace the foreign *CBF4* from grapevine.

**Figure 8.**
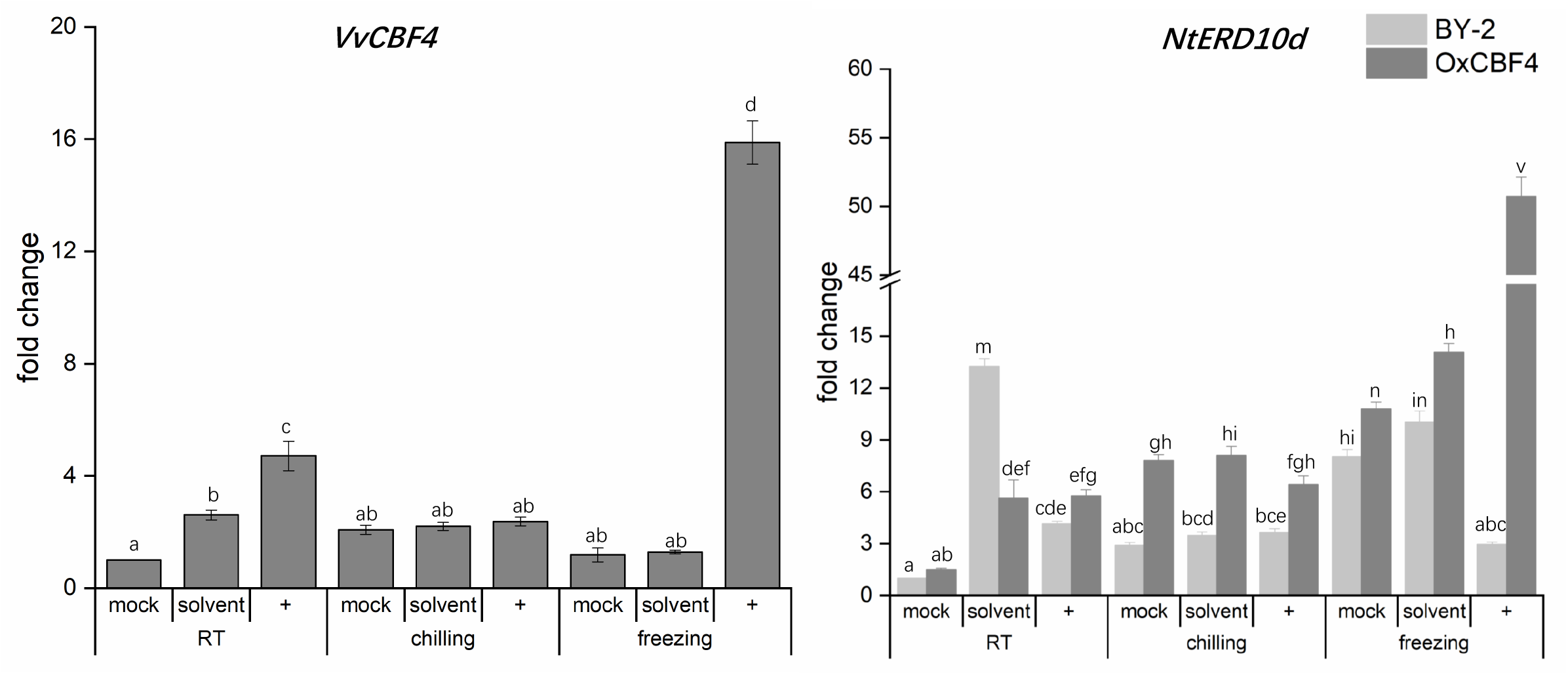
Effect of the lipoxygenase inhibitor phenidone on the expression of *VvCBF4* (in the VvCBF4ox line) and the COR transcript *NtERD10d* (in both, non-transformed tobacco BY-2 and the VvCBF4ox line) under 25°C (room temperature, RT), chilling (4°C), or freezing (0°C) stress. Expression was scored 24 h after the onset of the stress treatment either without phenidone (mock), or after pre-treatment for 30 min with 2 mM phenidone (represented by +) or same volume ethanol as solvent control. Data are normalised to the expression levels under room temperature without any of the inhibitors and represent means±SE of three independent experiments in technical triplicate. Statistically significant differences are represented as different letters in the LSD test (*P*<0.01). *NtERD10d* is based on its mRNA level in none-transformed BY-2 cells.

### Temporal cold-induced expression of CBF4 correlates in freezing resistance

To get insight into a possible link between CBF4 expression and cold tolerance, we followed CBF4 transcripts in leaves from different *Vitis* genotypes differing in their cold tolerance, either under chilling (4°C) or severe freezing (−20°C). Under chilling stress (**Fig. 9**), the cold tolerant *V. amurensis*from Northern China responded by a rapid (within 3 h) and persistent (over 12 h) accumulation of CBF4 transcripts. In contrast, *V. vinifera* Pinot Noir from Europe did not display any response of CBF4 to chilling. Interestingly, the cold sensitive *V. coignetiae* from Japan showed a significant elevation of *CBF4* transcript levels from the first time point but did not produce any significant induction during subsequent chilling stress. The patterns for severe freezing stress (−20°C) were significantly different. Here, *V. amurensis* accumulated *CBF4* transcripts slightly more sluggishly as compared to chilling stress, but again more persistently with a maximum at 12 h. In contrast, *V. vinifera* cv. Pinot Noir, which had not responded at all to chilling stress, produced a rapid, strong, but transient induction of *CBF4* transcripts peaking at 6 h and complete breakdown afterwards. The cold susceptible *V. coignetia* was not responsive at all in the beginning, and accumulated *CBF4* transcripts only from 12 h. Thus, the cold tolerance of *V. amurensis* correlated with a temporal pattern, where *CBF4* transcripts were induced not very swiftly (no response after 1 h of cold stress), but persistently (over 12 h or beyond), irrespective of the type of cold stress (chilling versus severe freezing). The intermediate *V. vinifera* Pinot Noir was not responsive to chilling stress, indicative of a poor cold acclimation, while it was not able to sustain expression under freezing stress. The susceptible *V. coignetia* was already challenged by chilling stress responding by a precocious induction but failed to respond appropriately under freezing stress. In addition to these genotype-dependency, the response pattern differed with respect to stress stringency.

**Figure 9.**
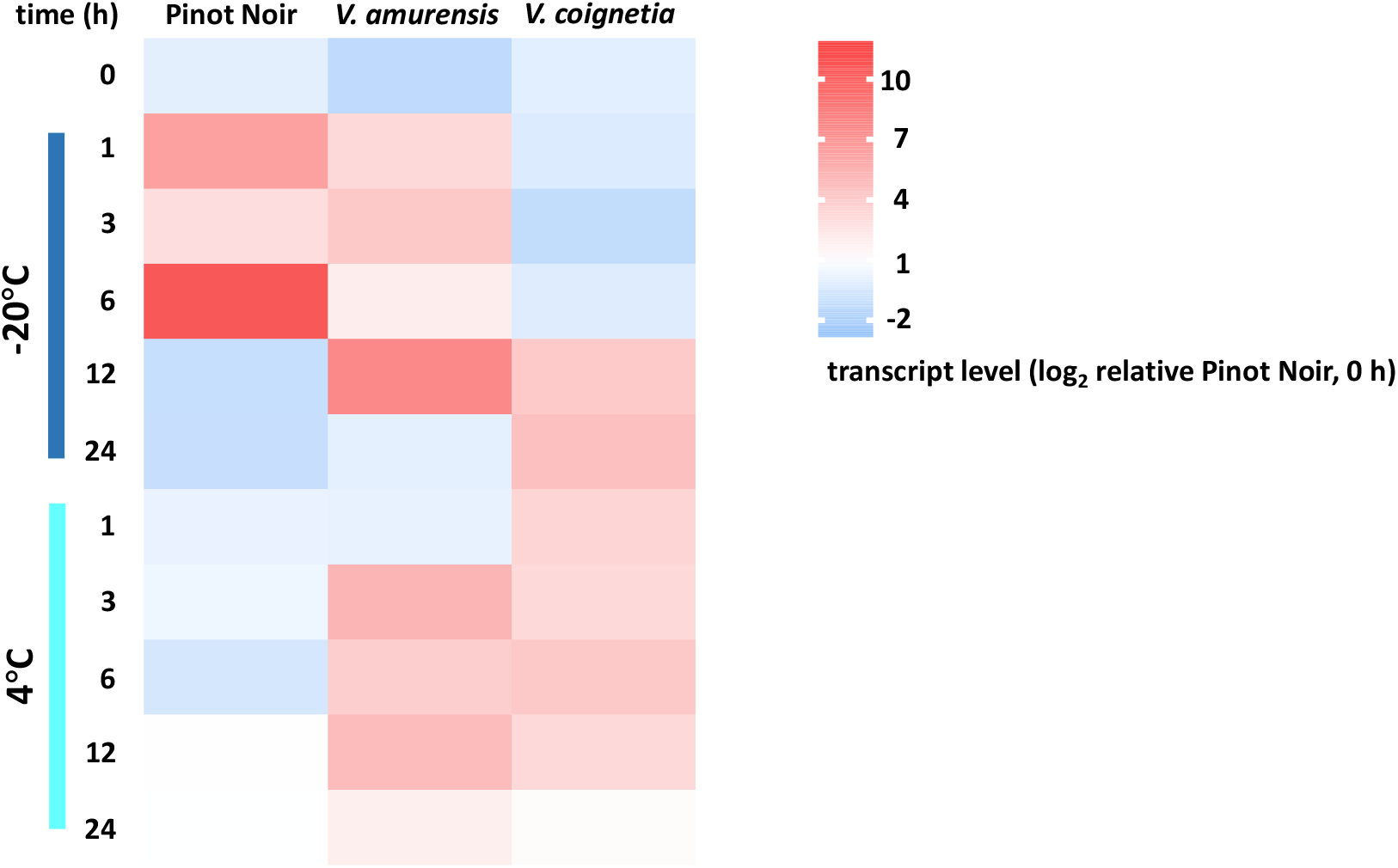
Time course for the response of transcripts for CBF4 to cold stress in fully expanded leaves from different *Vitis* genotypes. Plants of comparable developmental state were challenged by chilling (4°C) and freezing (−20°C). The leaves were sampled at the indicated time points after cold stress. Steady-state transcript levels are compared to the values found in Pinot Noir prior to cold stress. Data represent mean values from three biological replicates per each genotype and time point.

## Discussion

In addition to the usual three members of the CBF family, some species harbour a fourth member, CBF4, that is involved in cold acclimation, including important fruit crops such as Kiwi (Sun *et al*., 2021) or Garden Strawberry (Koehler *et al*., 2012). Also in grapevine, the expression of CBF4 correlates with cold tolerance (Xiao *et al*., 2008), and overexpression of CBF4 renders grapevines freezing tolerant (Tillett *et al*., 2012). To get more insight into the cellular mechanisms underlying the activity of grapevine CBF4 during chilling and freezing, we have generated tobacco cells overexpressing grapevine CBF4 as GFP fusion, and we followed the cold response of CBF4 in grapevine leaves from genotypes differing in their cold tolerance. In cell, chilling and freezing trigger distinct signalling pathway, including calcium influx, ROS burst, MAPK cascade, jasmonate and proteasome activity upstreaming of CBF transcript factors to specifically induce CBF4-dependent gene and cold responsive gene *ERD10d* depends on the quality of cold stress (chilling versus freezing).

These findings stimulate the following questions that are discussed below: By what mechanism is cold signalling translated into a change of gene expression? Which events are shared between chilling and freezing signals and where do they diverge? What is the function of CBF4 in the responses to chilling versus freezing? At what point do the CBF4 homologues from grapevine and tobacco differ and what might be the evolutionary context behind these differences?

### Nuclear import of CBF4 as a regulator, but not as the only regulator

In this study, the GFP-fusion of grapevine CBF4 is localised in the cytoplasm in the absence of cold stress but is seen in the nucleus when the expressing tobacco cells are exposed to 0°C (**Figure 1**). When we block nuclear import by GTP-γ-S, an inhibitor of Ran, the GTPase controlling nuclear import (Merkle *et al*., 1996), we block CBF4-dependent transcript responses, for instance of the transcription factors *NtDREB1* and *3*, as well as the endogenous *CBF4* orthologue *Avr9/Cf9,* and the cold responsive transcripts *NtERD10c* and *d*(**Figure 5**). These responses were qualitatively dependent on cold stress - under freezing stress, it is the transcription factors that were altered, under chilling stress, it was the *COR* transcripts.

To understand this contrasting pattern, it is important to link it with the expression of these transcripts between the non-transformed wildtype and VvCBF4 overexpressor (**Figure 4**). Here, the accumulation of the *COR* transcripts *NtERD10c* and *d* in response to chilling was clearly suppressed, when the import of CBF4 was blocked by GTP-γ-S, meaning that *VvCBF4* is suppressing accumulation of the COR transcripts under chilling. Interestingly, there is no effect of GTP-γ-S on the response of these transcripts to freezing (**Figure 5**). Instead, these transcripts are more efficiently induced under freezing, if CBF4 is overexpressed (**Figure 4**).

In contrast, the response of the transcription factors is more sensitive to GTP-γ-S under freezing stress, whereby the tobacco CBF4 homologues *Avr9/Cf9* and *DREB3* are inhibited, while DREB1 is stimulated, when nuclear transport is blocked (**Figure 5**). The amplitude of the DREB3 response to freezing is enhanced under freezing stress (**Figure 4**), and, thus, parallels the pattern seen for *ERD10c* and *d*. The endogenous *CBF4 (Avr9/Cf9)* is induced independently of, whether VvCBF4 is present or not (**Figure 3C, D**).

The most straightforward model to explain these results is that CBF4, upon nuclear import in response to chilling, is suppressing accumulation of the COR transcripts *ERD10c* and *d*. In response to freezing, the import of CBF4 stimulates the accumulation of *DREB3* and *Avr9/Cf9,* while suppressing that of *DREB1.* The nuclear import of CBF4 has, thus, a different effect, depending on the temperature of the cold stress. Such a sign reversal cannot be explained by a model where nuclear import of CBF4 is the only switch. There must be a second factor that acts in concert with CBF4 and is differentially activated or deployed, depending on the severity of cold stress.

To find a transcription factor in the cytoplasm, may seem unexpected at first glance. In fact, in those cases, where the subcellular localisation of CBFs from type 1-3 was addressed, usually by ectopic expression of GFP fusions in onion epidermal cells, *N benthamiana* leaves, or in protoplasts, they were seen in the nucleus (Aloe, (Wang and He, 2007); Tea Plant, (Hu *et al*., 2020), *Deschampsia antarctica,* (Byun *et al*., 2015); *Phyllostachys edulis,* (Liu *et al*., 2012); Soybean, (Kidokoro *et al*., 2015). The activity of these factors does not seem to be brought about by their import. Instead, it is the import of other activating factors that unlocks CBF activity. For instance, specific cytosolic thioredoxins respond to cold-induced oxidative burst at the plasma membrane by nuclear import and cleave their sulphur bonds that had sequestered CBF oligomers that are then released and trigger the expression of COR genes (Wi *et al*., 2022).

Is shuttling of a transcription factor itself a specific feature delineating CBF4s, from the other CBFs? The different mode of activation might be linked with their specific role in cold acclimation. The fact that even in *Vitis* itself a CBF1-GFP fusion is exclusively seen in the nucleus (Xiao *et al*., 2006) would support such a model, where the difference in function is reflected as difference in activation mechanisms. It should be mentioned, however, that the CBF4 from *Arabidopsis* seems to differ from *VvCBF4.* Here, a CBF4-YFP fusion was found preferentially in nuclei upon transient expression in *N benthamiana* leaves and *Arabidopsis* hypocotyls (Vonapartis *et al*., 2022), albeit the interpretation of the shown images is not conclusive, because there is cytoplasmic background and the resolution of those images does not suffice to detect a perinuclear localisation (which was also not in the focus of that work).

Regulation of gene expression by signal-dependent import of transcription factors is a phenomenon that is well known from animals. A classic example is the glucocorticoid receptor that is bound to heat-shock protein 90 and a tetratricopeptide repeat containing protein, but detaches upon binding of steroid hormones, enters the nucleus, and acts there as transcriptional regulator (for review see (Echeverría *et al*., 2009)). Also, for plants, evidence for signal-dependent import of transcriptional regulators as common regulatory paradigm has accumulated. Activation of the plant photoreceptor phytochrome will release the bZIP transcription factor CPRF2 from its cytoplasmic tether, shifting it from the cytosol into the nucleus (Kircher *et al*., 1999). Also, cold stress can send transcription factors to the nucleus as found for *VaWRKY12*a transcription factor from the Siberian wild grapevine *V. amurensis*that repartitions from the cytoplasm to the nucleus in response to low temperature (Zhang *et al*., 2019). Recently, an unconventional class-XIV kinesin motor, Dual Localisation Kinesin, was found to shuttle to the nucleus, when the microtubules disassemble in the cold (Xu *et al*., 2018). In the nucleus, this kinesin binds specifically to a motif in the promoter of CBF4 and modulates its expression (Xu *et al*., 2022). Thus, our finding that nuclear import of CBF4 acts as regulator for cold-dependent transcription adds another example for this type of regulatory paradigm.

### Chilling and freezing signalling differ in the role of oxidative burst and of jasmonate

A part of the complexity and even inconsistency about the role of different events in cold signalling seems to arise because molecular genetics of the chilling insensitive model *Arabidopsis thaliana* and the chilling sensitive model rice are often listed together (for a critical discussion of this problem see (Wang *et al*., 2020)). We have, therefore, used the approach to compare signalling evoked by chilling and freezing stress in the same model, tobacco BY-2 cells. By this approach we are able to see that the dependence of cold-induced transcripts on different signalling events depends on the type of cold stress (**Fig. 7**). In the following, we want to highlight two of these events:

Using *NtERD10d* as proxy for COR gene expression, we observed that the solvent DMSO, which is also a membrane rigidifier, was enhancing the induction of this COR transcript under cold stress. Since DMSO evokes a drop of membrane fluidity and, thus, mimicks cold, the stimulating effect of DMSO under cold stress is straightforward to understand (Sangwan *et al*., 2001; Wang and Nick, 2017). This stimulation is completely (chilling) or partially (freezing) eliminated by Diphenylene Iodonium (DPI), which in plants is a specific inhibitor of the membrane-located NADPH oxidase Respiratory burst oxidase Homologue (for a discussion of this inhibitor see (Eggenberger *et al*., 2017)). Thus, an oxidative burst occurring at the plasma membrane is required for the cold-induced induction of COR transcripts happening in the nucleus. A molecular candidate for this transduction event might be thioredoxins such as AthTrx-h2 that translocate to the nucleus in response to an oxidative burst, unleashing there CBFs from sulphur bridge bondage (Wi *et al*., 2022). A (non-intuitive) implication of this model would be that overexpression of *VvCBF4* would stimulate the import of CBF4 under freezing stress amplifying the cold response of *ERD10d*transcripts (**Fig. 4**), while under chilling, the accumulation of these transcripts should remain sensitive to DPI. This is exactly, what we observe: For the VvCBF4 overexpressor, the inhibition by DPI was seen only for chilling stress, while for freezing stress, DMSO was not inducive, and DPI (dissolved in DMSO) was giving the same transcript level as the mock control.

A second difference in chilling and freezing signalling was seen for the dependence on phenidone, inhibiting the conversion of α-linolenic acid into 13-HPOT, the first committed metabolite of the jasmonate pathway (Bruinsma *et al*., 2010a). Here, we see a very strong upregulation of *ERD10d* and of *CBF4* transcripts in the CBF4 overexpressor, while this induction of COR transcripts is not observed in the WT (**Fig. 8**). Moreover, this phenomenon is only seen for freezing, not for chilling stress. Thus, jasmonates are negative regulators of *CBF4* expression and CBF4-dependent COR-expression under freezing stress (**Fig. 8**). A mild induction of *CBF4* and of *ERD10d* transcripts was also seen for room temperature, suggesting that the phenomenon is under control of a process that acts also in the absence of cold stress. A prime candidate is ICE1, which is continuously synthetised *de novo* and degraded in the warm (Chinnusamy *et al*., 2007), but accumulates, when proteolysis is inhibited by post-translational modifications (Guo *et al*., 2018). JA-Ile induces expression of the JAZ response regulators, specific members (in *Arabidopsis thaliana,* JAZ1 and JAZ4) repress the binding of ICE1 to the promoters of its target genes (including the CBFs) and, thus, downmodulates cold-dependent gene expression (Hu *et al*., 2013). An implication of such a mechanism would be that phenidone should upregulate the expression of *CBF4* and of CBF4-dependent COR expression (*ERD10d*). This is, what we observe (**Fig. 8**). A further implication of this model is that inhibition of proteasome activity should upregulate both transcripts. Also, this implication was tested by us and confirmed by using MG132 (**Fig. 7**). Moreover, the elimination of all responses by Actinomycin shows that de-novo protein synthesis is needed, lending further support to this model (**Fig. 6**). A third implication of the model would be that overexpression of CBF4 should override this mechanism, because ICE1 is not required. Again, we did not observe the upregulation of CBF4 under freezing (**Fig. 7**), while *ERD10d* transcripts were induced, indicating that, here, the effect of the other CBFs that are controlled by ICE1, kicks in. The fact that MG132 remains effective under chilling stress (**Fig. 7**), provides a further example that signalling depends on the quality of cold stress.

We arrive at a working model, supported by numerous, non-intuitive implications that could be experimentally tested and confirmed, where chilling and freezing stress are transduced by two concurrent signalling chains, whose relative contribution depends on the stringency of stress. One chain works through nuclear import of CBF4 and is not dependent on oxidative burst at the plasma membrane, the second chain works through a signal deployed by oxidative burst and is independent of CBF4. Under freezing stress, CBF4-dependent signalling dominates, under chilling stress, the concurrent pathway is preponderant. Whether this second pathway employs thioredoxins (Wi *et al*., 2022) remains to be elucidated in future studies.

### Survival under freezing: it is the response of CBF4 to chilling that matters

Transcripts for the introduced *VvCBF4* were strongly induced by chilling, barely by freezing stress (**Fig. 3**). The mitigating effect on mortality was, however, seen for freezing stress in the first place (**Fig. 2**), because chilling did not lead to substantial mortality, indicating that tobacco BY-2 cells are fairly chilling tolerant. In congruence with this notion, the transcripts of the endogenous tobacco *Avr9/Cf9* produced only a small increased in response to chilling. Thus, the behaviour of CBF4 reflected the level of cold tolerance of the respective progenitor species. Since the expression of *VvCBF4* was driven by the constitutive CaMV 35S, the elevated mRNA level of *VvCBF4* in the transgenic line (**Fig. 3**) was to be expected. However, the fact that this transcript level increased even further under chilling, must be due to post-transcriptional regulation. In other words, transcript stability must be under control of cold-induced signalling, which was also reported for other CBFs (Hwarari *et al*., 2022).

To get further insight into the physiological context of CBF4 accumulation, we addressed transcript regulation during chilling and freezing in three grapevine species differing in their cold tolerance (**Fig. 9**). Interestingly, it is not the induction of CBF4 under freezing stress - this is far more pronounced in Pinot Noir. In fact, it is the rapid induction of CBF4 during chilling that can serve as hallmark for superior cold tolerance. In other words, the tough fellows are those that are the most sensitive to chilling. A previous study arrived at a similar conclusion comparing the responses of winter-wheat varieties differing in freezing tolerance (Abdrakhamanova *et al*., 2003). Also in that study, a swift and sensitive response to chilling was correlated with the ability to deploy a high degree of freezing tolerance.

It is not low temperature as such that renders blurred seasonality such a big problem in agriculture, it is the suddenness of temperature drops that lead to damage. A species adapted to the continental conditions of Mandzhuria (*V. amurensis)* has evolved a very efficient signalling leading to a swift induction of CBF4, such that cold acclimation can be deployed rapidly and efficiently as prerequisite to survive in an environment, where drastic increases and drops of temperature are common. In contrast, *V. coignetia*, which comes from a much milder and temperate climate, was not selected for such a swift induction. Likewise, *V. vinifera* that had survived pleistocenic glaciation in refugia in the Mediterranean and the Black Sea, while able to accumulate CBF4 under freezing, is not very responsive to the chilling that under natural conditions precedes a freezing episode.

These evolutionary considerations shift the upstream signalling driving the induction of CBF4 into the focus of interest. Is it the steady-state levels of signalling compounds, such as RboH, is it amplifications of the primary sensory events at the plasma membrane that account for the strong chilling response in *V. amurensis?* Is it a second, unknown factor that accompanies nuclear import of CBF4 and modifies the transcriptional response as suggested by the differences in the response pattern for chilling and freezing in the CBF4 overexpressor. The recent finding that the unconventional class-XIV kinesin Dual Localisation Kinesin, in response to cold stress, can shuttle from the membrane to the nucleus and specifically modulate the expression of the tobacco CBF4 homologue *Avr9/Cf9* (Xu *et al*., 2022) opens new exciting questions on this second signalling pathway and its interaction with CBF4 dependent cold responses.

## Acknowledgements

We thank Sabine Purper, Gabi Jürges for the assistance and our gardener Kevin Malakowsky for looking after the grapevine plants. This work was supported by a fellowship from the China Scholarship Council to Wenjing Shi.

